# Professional Swimmers and Normal Populations have Different Patterns of Epidermal Green Autofluorescence

**DOI:** 10.1101/510891

**Authors:** Yue Tao, Mingchao Zhang, Yujia Li, Xunzhang Shen, Weihai Ying

## Abstract

Our recent studies have suggested that green autofluorescence of the fingernails and certain regions of skin may be novel biomarkers for disease diagnosis and evaluation of the injury state of cardiovascular system. Our study has also suggested that oxidative stress may produce the increased epidermal green AF by altering keratin 1. Since athletes’ body may have increased oxidative stress and inflammation, we proposed our hypothesis that athletes may have increased green AF in their fingernails and certain regions of their skin. In current study we tested this hypothesis by determining the green AF of professional swimmers. We found that compared with age-matched controls, both the green AF intensity and AF asymmetry in both right and left antebrachium and Ventriantebrachium of the professional swimmers is significantly higher. In left Dorsal Centremetacarpus, the green AF intensity of the professional swimmers is significantly higher than that of the age-matched controls. In contrast, the green AF intensity or AF asymmetry of the professional swimmers is not significantly different from that of the age-matched controls in Centremetacarpus, Ventroforefinger, Dorsal Index Finger and Index Fingernails. Collectively, our study has provided first evidence suggesting that athletes have increased green AF intensity and asymmetry in certain regions of their skin. Based on this finding, we may evaluate non-invasively the levels of oxidative damage and inflammation in athletes’ body.

## Introduction

It is established that regular exercise can produce beneficial effects on physiological adaptations in the body. However, intense and prolonged physical exercises may also produce detrimental effect on the body. A number of studies have indicated that intense physical exercises can lead to increased oxidative stress (9,12,13), which may be caused by such factors as mitochondrial electron leakage and NADPH oxidase activity (13). It has also been reported that intense physical exercises can lead to increased inflammation in the body (9–11). In order to prevent athletes from damage produced by intense exercise-induced oxidative stress and inflammation, it is necessary to develop non-invasive approaches to monitor the oxidative stress and inflammation in athletes’ body. However, so far virtually all of the approaches used for determining oxidative stress and inflammation in the body require blood drawing.

Human autofluorescence (AF) has been used for non-invasive diagnosis of such diseases as diabetes (3,8). Our recent studies have suggested that characteristic ‘Pattern of Epidermal Green Autofluorescence (AF)’ of each disease may be novel biomarkers for non-invasive diagnosis of a number of diseases, including acute ischemic stroke (AIS) (1), myocardial infarction (MI) (14), stable coronary artery disease (14), Parkinson’s disease (2) and lung cancer (7). Our study has also shown that the AF intensity in certain regions of skin is highly correlated with a person’s risk to develop AIS, suggesting that the AF may also become a biomarker for evaluation of injury levels of vascular system (6).

Our recent study has suggested that oxidative stress is an important factor that can increase the epidermal green AF by altering keratin 1 (4,5). Our latest study has also indicated that inflammation can induce increased green AF of mouse’s skin (15). Based on these findings, we proposed our hypothesis that athletes may have increased epidermal green AF in certain regions of their skin, which may be produced by the increased oxidative stress and inflammation in their body. In our current study, we tested this hypothesis by determining the green AF of professional swimmers and age-matched controls. Our study has indicated that in the Antebrachium and Ventriantebrachium, professional swimmers have significant higher levels of epidermal green AF intensity and asymmetry.

## Methods

### Human subjects

The human subjects in our study were divided into two groups. Group 1: Professional swimmers; and Group 2: Age-matched controls. The age of all of the subjects is 18 years of old.

### Autofluorescence determinations

For all of the human subjects, the AF intensity in the following seven regions on both hands, i.e., fourteen regions in total, was determined, including the index fingernails, Ventroforefingers, dorsal index fingers, Centremetacarpus, Dorsal Centremetacarpus Ventriantebrachium, and Dorsal Antebrachium. A portable AF imaging equipment was used to detect the AF of the these fourteen regions of the human subjects. The excitation wavelength is 485 nm, and the emission wavelength is 500 - 550 nm.

### Statistical analyses

All data are presented as mean + SEM. Data were assessed by student ŕ-test, except where noted. *P* values less than 0.05 were considered statistically significant.

## Results

We determined the green AF intensity in multiple regions of the skin of the professional swimmers and the age-matched controls. We found that compared with the age-matched controls, both the green AF intensity and AF asymmetry in both right and left Antebrachium of the professional swimmers is significantly higher (Figs. 1A and 1B). Similarly, in Ventriantebrachium, both the green AF intensity and AF asymmetry of the professional swimmers is significantly higher than that of the age-matched controls (Figs. 2A and 2B).

**Fig. 1.**
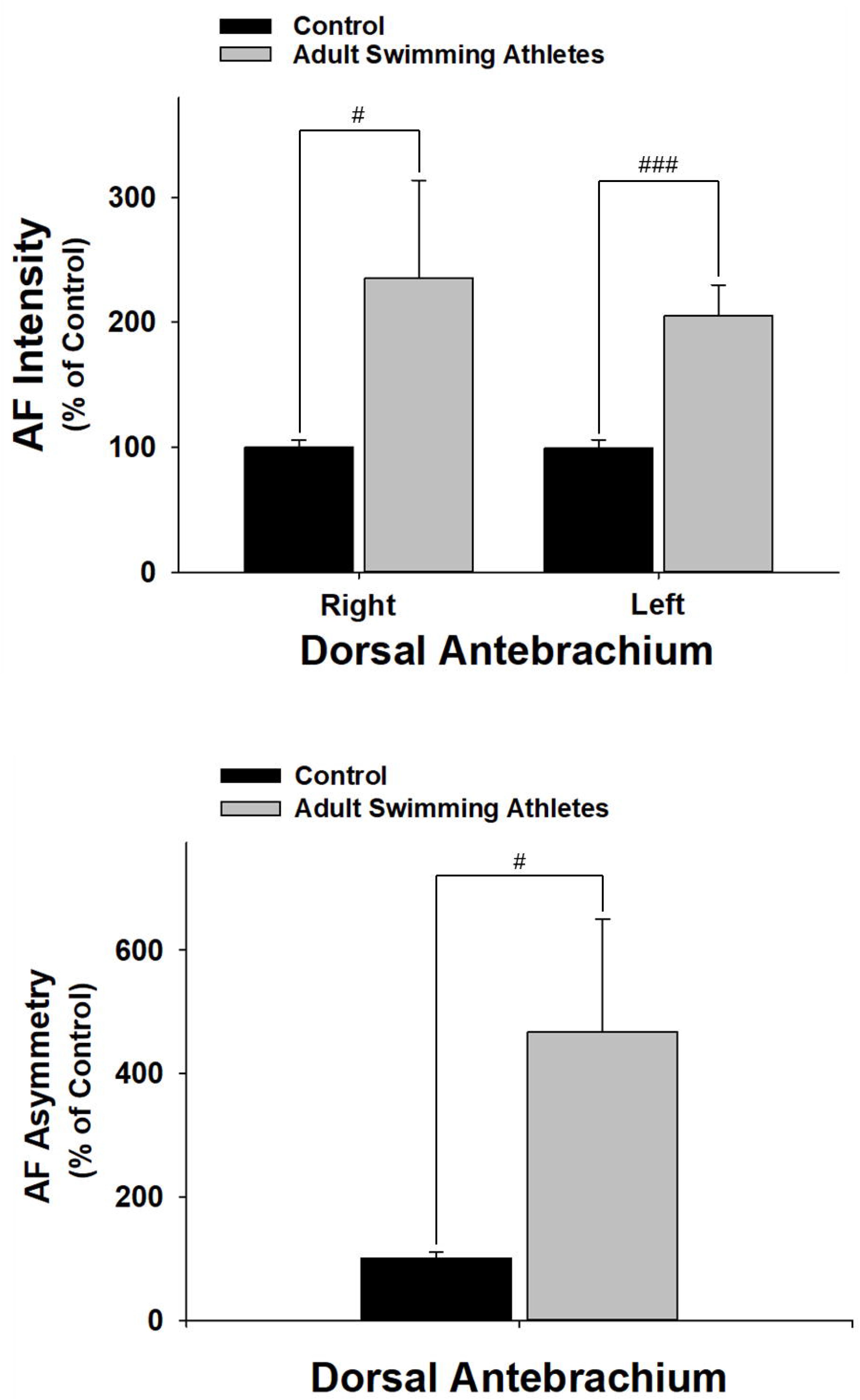
Both the green AF intensity and AF asymmetry in both right and left Antebrachium of the professional swimmers is significantly higher that of the age-matched controls. The number of the age-matched controls and the professional swimmers is 50 and 32, respectively. #, *p* < 0.05; ###, *p* < 0.001 (Student *t*-test).

**Fig. 2.**
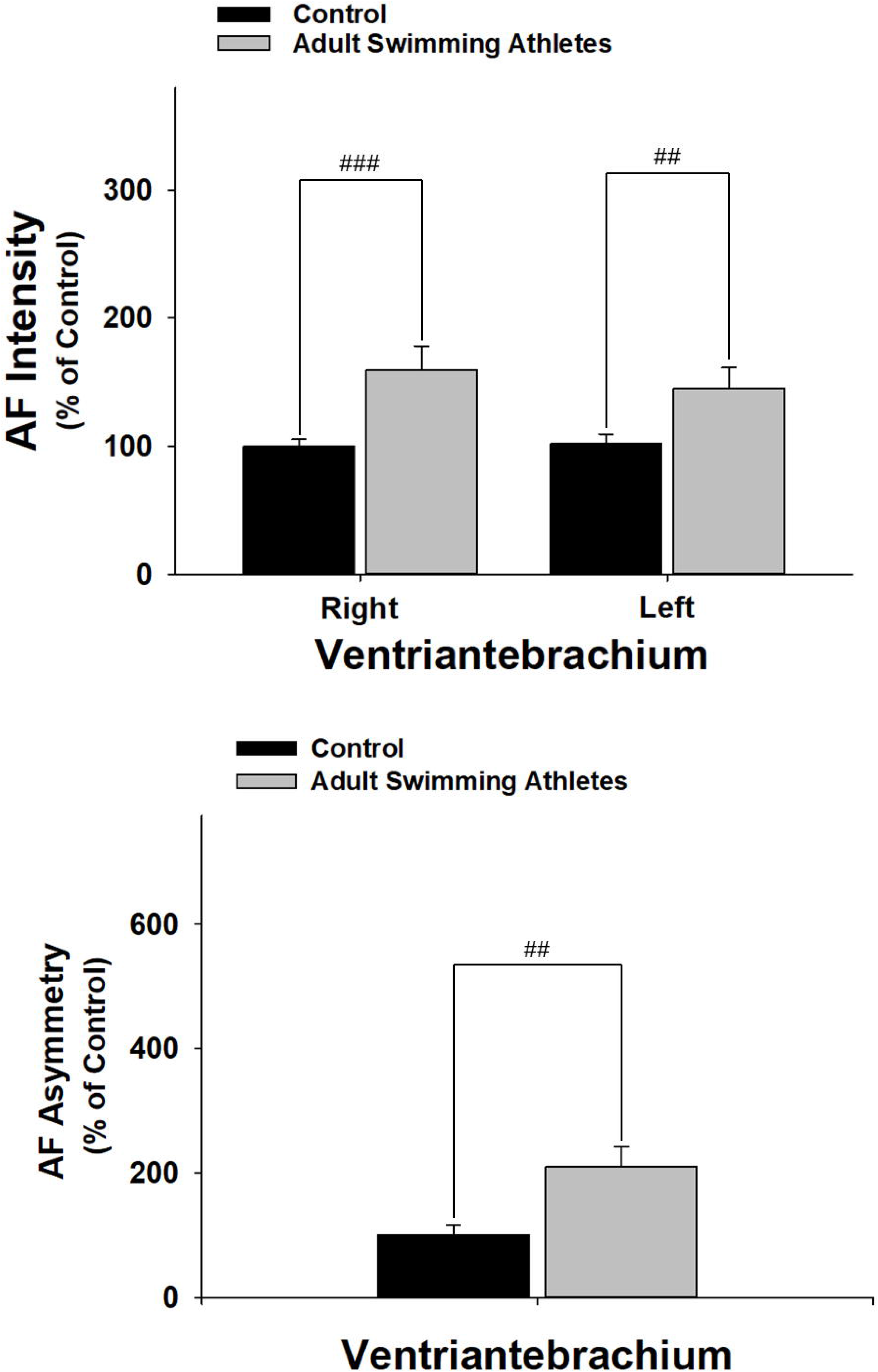
Both the green AF intensity and AF asymmetry in both right and left Ventriantebrachium of the professional swimmers is significantly higher than that of the age-matched controls. The number of the age-matched controls and the professional swimmers is 50 and 32, respectively. ##, *p* < 0.01; ###, *p* < 0.001 (Student *t*-test).

In left Dorsal Centremetacarpus, the green AF intensity of the professional swimmers is significantly higher than that of the age-matched controls (Figs. 3A and 3B). In contrast, in Centremetacarpus (Figs. 4A and 4B), Ventroforefinger (Figs. 5A and 5B), Dorsal Index Finger (Figs. 6A and 6B) and Index Fingernails (Figs. 7A and 7B), the green AF intensity or AF asymmetry of the professional swimmers is not significantly different from that of the age-matched controls.

**Fig. 3.**
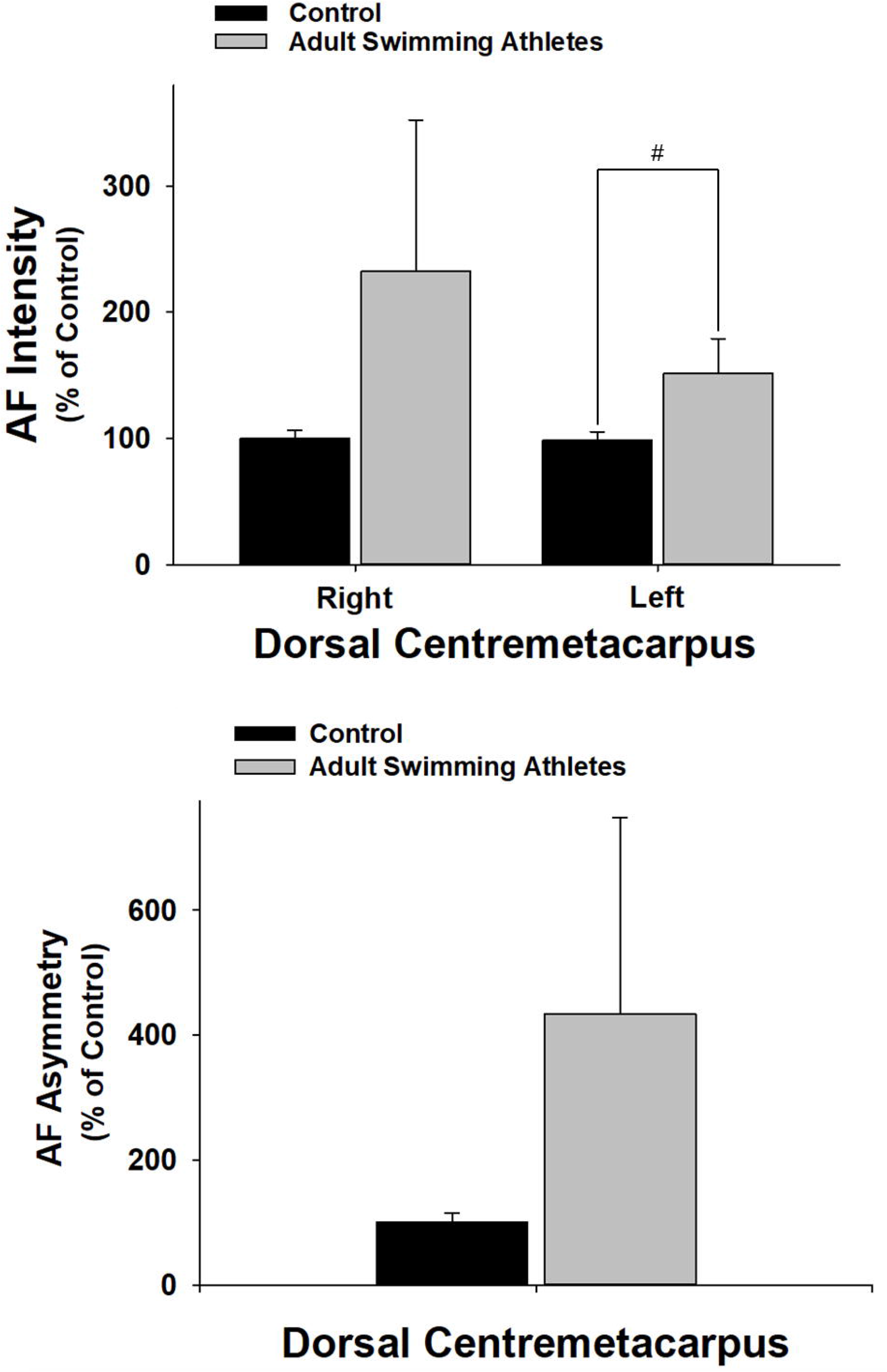
The green AF intensity of the professional swimmers is significantly higher than that of the age-matched controls in left Dorsal Centremetacarpus. The number of the age-matched controls and the professional swimmers is 50 and 32, respectively. #, *p* < 0.05 (Student *t*-test).

**Fig. 4.**
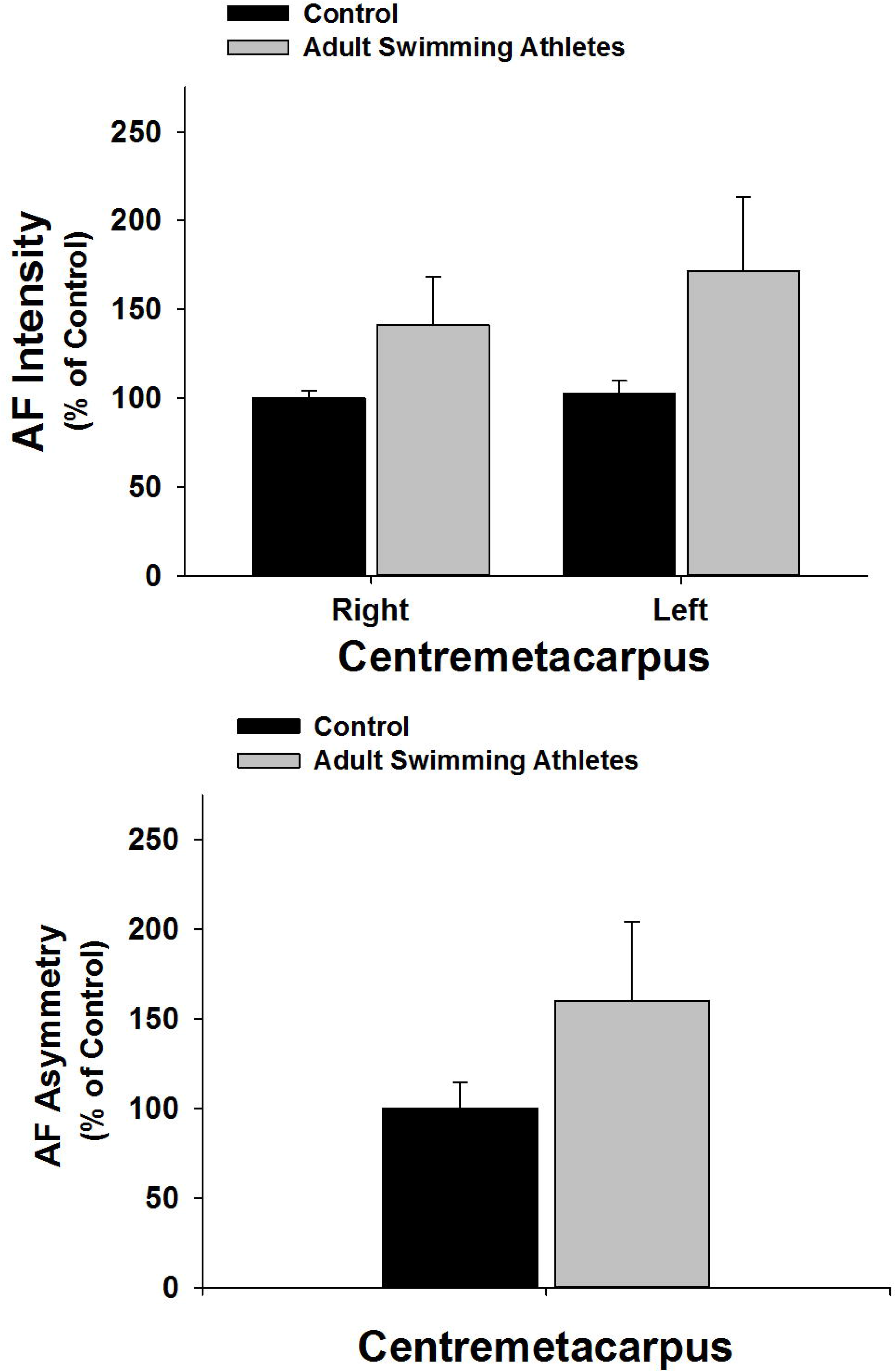
In Centremetacarpus, the green AF intensity or AF asymmetry of the professional swimmers is not significantly different from that of the age-matched controls. The number of the age-matched controls and the professional swimmers is 50 and 32, respectively.

**Fig. 5.**
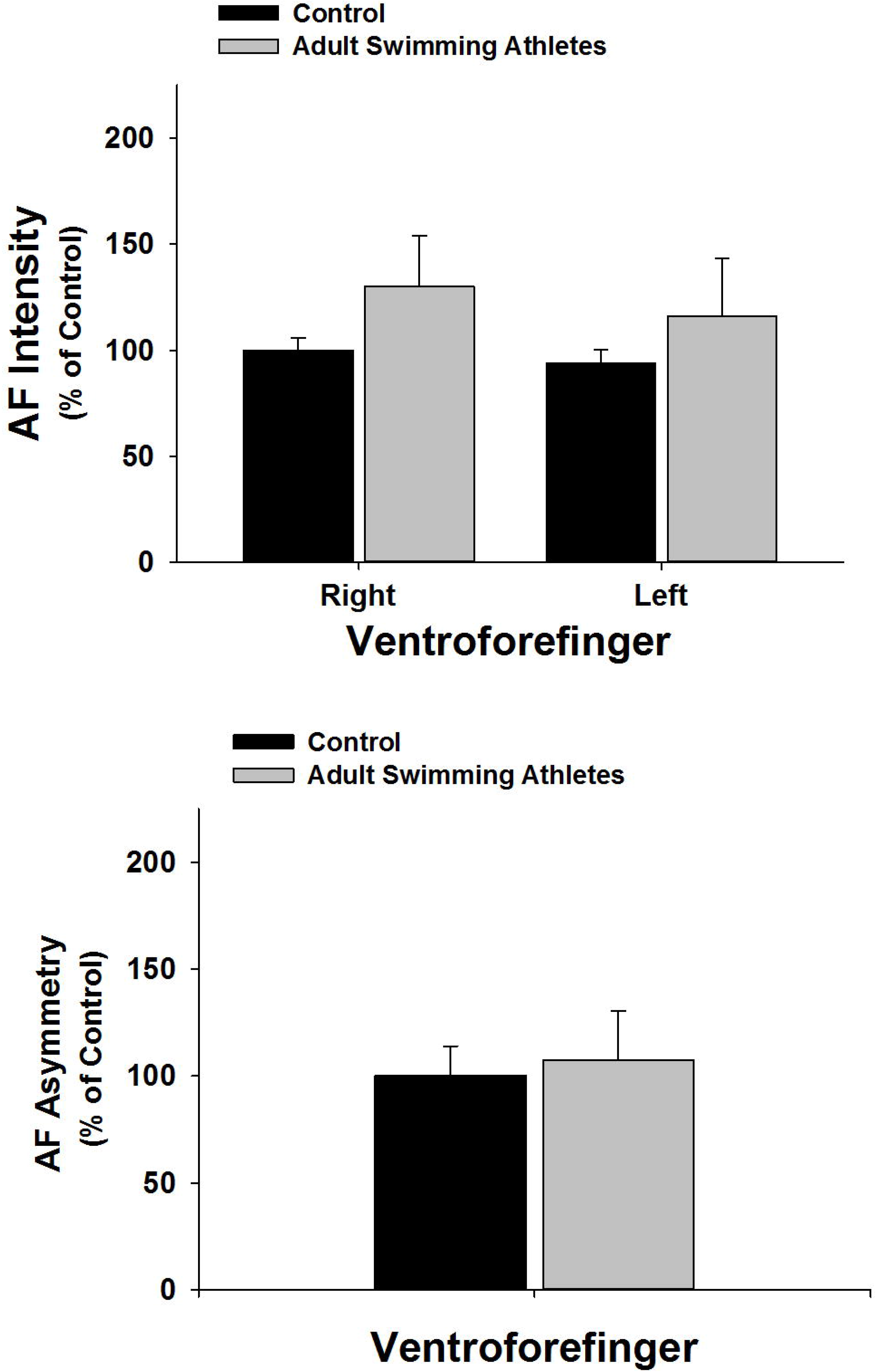
In Ventroforefinger, the green AF intensity or AF asymmetry of the professional swimmers is not significantly different from that of the age-matched controls. The number of the age-matched controls and the professional swimmers is 50 and 32, respectively.

**Fig. 6.**
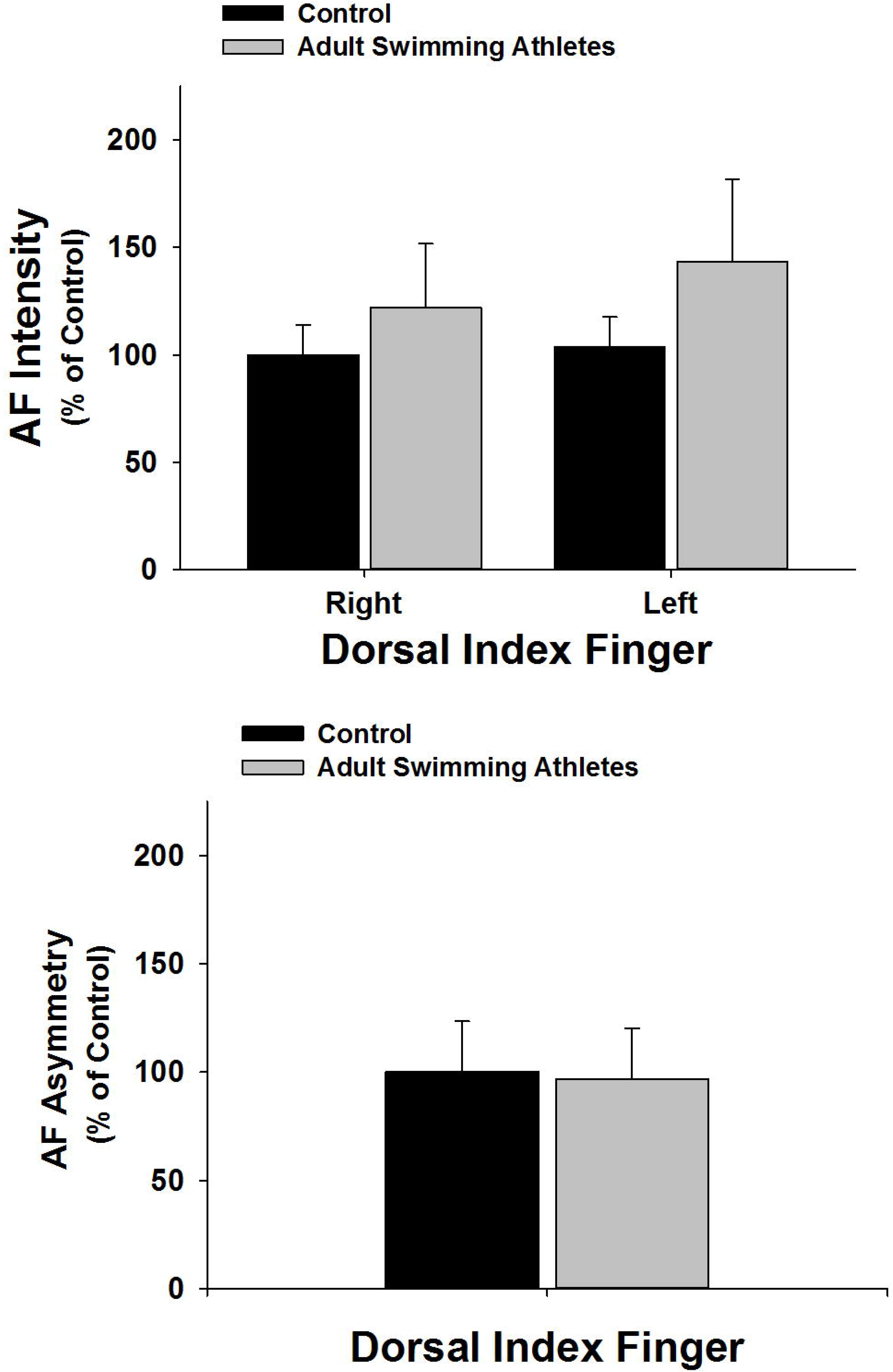
In Dorsal Index Finger, the green AF intensity or AF asymmetry of the professional swimmers is not significantly different from that of the age-matched controls. The number of the age-matched controls and the professional swimmers is 50 and 32, respectively.

**Fig. 7.**
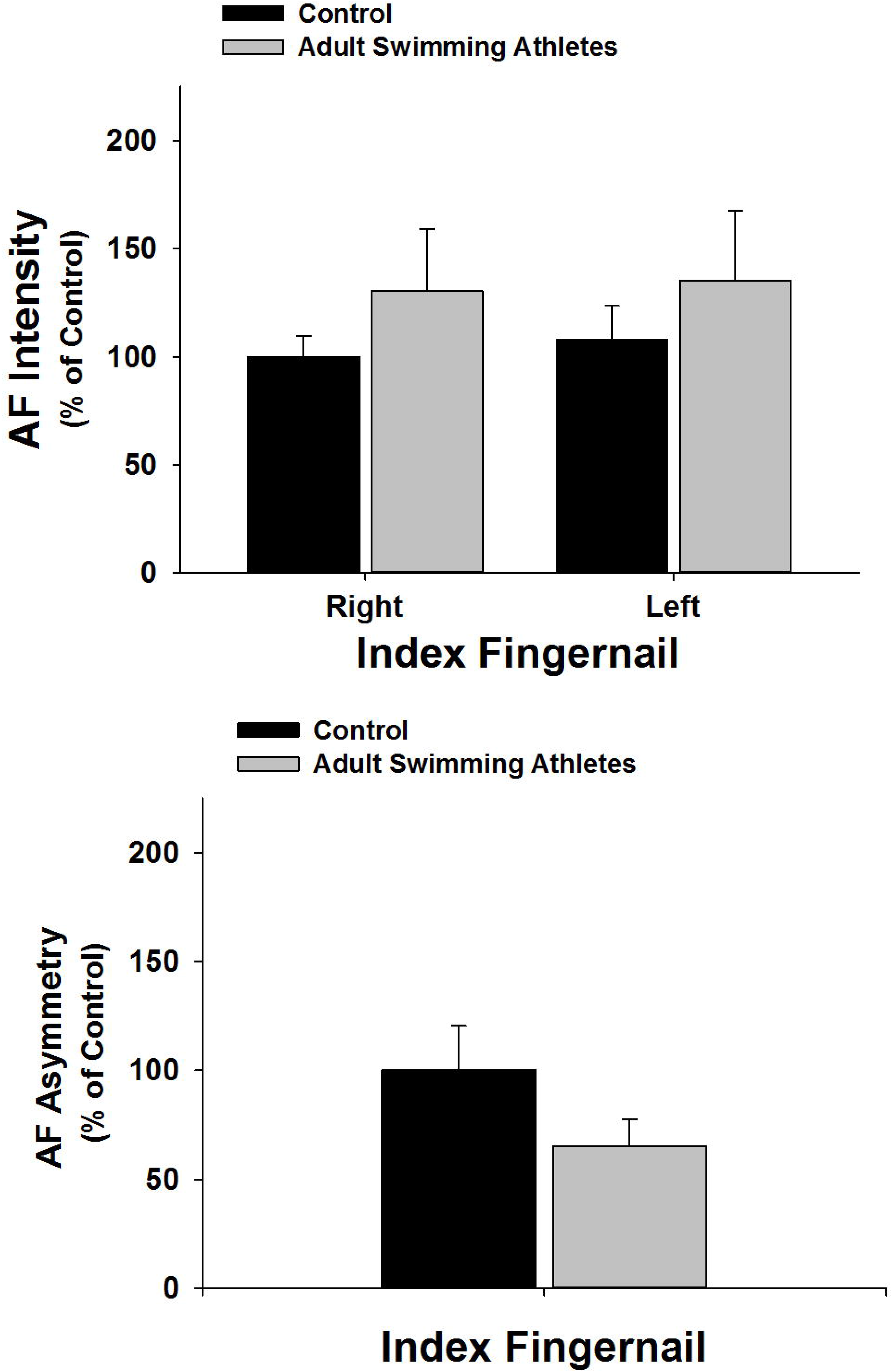
In Index Fingernails, the green AF intensity or AF asymmetry of the professional swimmers is not significantly different from that of the age-matched controls. The number of the age-matched controls and the professional swimmers is 50 and 32, respectively.

## Discussion

The major findings of our current study include: First, compared with the age-matched controls, both the green AF intensity and AF asymmetry in both right and left Antebrachium of the professional swimmers is significantly higher; second, both the green AF intensity and AF asymmetry of the professional swimmers is significantly higher than that of the age-matched controls in Ventriantebrachium; third, the green AF intensity of the professional swimmers is significantly higher than that of the age-matched controls in left Dorsal Centremetacarpus; and fourth, the green AF intensity or AF asymmetry of the professional swimmers is not significantly different from that of the age-matched controls in Centremetacarpus, Ventroforefinger, Dorsal Index Finger and Index Fingernails.

It has been indicated that intense physical exercises can lead to increased oxidative stress (9,12,13). Multiple studies have also reported that intense physical exercises can lead to increased inflammation in the body (9–11). Therefore, it is valuable to develop non-invasive approaches to monitor the oxidative stress and inflammation in athletes’ body, which may prevent athletes from damage produced by intense exercise-induced oxidative stress and inflammation. Our recent study has suggested that oxidative stress is an important factor that can increase the epidermal green AF by altering keratin 1 (4,5). Our latest study has also indicated that inflammation can induce increased green AF of mouse’s skin (15). Based on these observations, we investigated the possibility that professional swimmers may have increased epidermal green AF. Our findings have suggested that professional swimmers have significantly increased epidermal green AF intensity and asymmetry in both Antebrachium and Ventriantebrachium. These findings may establish a critical basis for non-invasive monitoring of the levels of oxidative stress and inflammation in athletes’ body.

Our study has found that professional swimmers have increased epidermal green AF asymmetry in both Antebrachium and Ventriantebrachium. Our previous studies have also found that the green AF of the patients of such diseases as AIS (1) and MI (14) is asymmetrical in several regions of their skin. Future studies are warranted to elucidate the mechanism underlying the asymmetry of the epidermal AF intensity of these populations.

Our study has found that the AF intensity is increased selectively in certain regions of the skin of professional swimmers, including both right and left Antebrachium and Ventriantebrachium as well as left Dorsal Centremetacarpus. We proposed that the selective AF increases may indicate the selective increases in the oxidative stress and/or inflammation of certain tissues and organs which are associated with those regions of skin. Future studies are necessary to expose the mechanisms underlying these observations. Moreover, it is necessary to determine if the athletes of various types of sports have similar patterns of AF.

## Acknowledgment

The authors would like to acknowledge the financial support by a Major Special Program Grant of Shanghai Municipality (Grant # 2017SHZDZX01) (to W.Y.), and a Major Research Grant from the Scientific Committee of Shanghai Municipality #16JC1400500 and #16JC1400502 (to W.Y. and X.S.).

## References

1. Wu D, Zhang M, Tao Y, Li Y, Zhang S, Chen X, Ying W. Asymmetric increases in the intensity of the green autofluorescence of ischemic stroke patients’ skin and fingernails: A novel diagnostic biomarker for ischemic stroke. bioRxiv 310904, 2018.

4. Wu D, Zhang M, Tao Y, Li Y, Shen L, Li Y, Ying W. Selectively increased autofluorescence at fingernails and certain regions of skin: A potential novel diagnostic biomarker for Parkinson disease. bioRxiv 322222, 2018.

3. Meerwaldt R, Links T, Graaff R, Thorpe SR, Baynes JW, Hartog J, Gans R, Smit A. Simple noninvasive measurement of skin autofluorescence. Ann N Y Acad Sci 1043: 290–8, 2005.

4. Zhang M, Dhruba TM, He H, Li Y, Yan W, Yan W, Zhu Y, Ying W. UV-Induced Keratin 1 Proteolysis Mediates UV-Induced Skin Damage. bioRxiv 226308, 2018.

11. Zhang M, Dhruba TM, Li Y, Ying W. Oxidative stress mediates UVC-induced increases in epidermal autofluorescence of C57 mouse ears. BioRxiv 298000, 2018.

12. Zhang M, Wu D, Tao Y, Li Y, Zhang S, Chen X, Ying W. Green autofluorescence intensity of skin and fingernails: A novel biomarker for non-invasive evaluation of pathological state of blood vessels. bioRxiv 403832, 2018.

13. Zhang M, Tao Y, Chang Q, Li Y, Chu T, Ying W. Selectively increased autofluorescence at certain regions of skin may become a novel diagnostic biomarker for lung cancer. bioRxiv 315440, 2018.

8. Moran C, Munch G, Forbes JM, Beare R, Blizzard L, Venn AJ, Phan TG, Chen J, Srikanth V. Type 2 diabetes, skin autofluorescence, and brain atrophy. Diabetes 64: 279–83, 2015.

9. Nogueira JE, Passaglia P, Mota CMD, Santos BM, Batalhao ME, Carnio EC, Branco LGS. Molecular hydrogen reduces acute exercise-induced inflammatory and oxidative stress status. Free Radic Biol Med 129: 186–193, 2018.

10. Peake JM, Della Gatta P, Suzuki K, Nieman DC. Cytokine expression and secretion by skeletal muscle cells: regulatory mechanisms and exercise effects. Exerc Immunol Rev 21: 8–25, 2015.

11. Pedersen BK, Hoffman-Goetz L. Exercise and the immune system: regulation, integration, and adaptation. Physiol Rev 80: 1055–81, 2000.

12. Powers SK, Radak Z, Ji LL. Exercise-induced oxidative stress: past, present and future. J Physiol 594: 5081–92, 2016.

13. Rahal A, Kumar A, Singh V, Yadav B, Tiwari R, Chakraborty S, Dhama K. Oxidative stress, prooxidants, and antioxidants: the interplay. Biomed Res Int 2014:761–264, 2014.

14. Qu X, Li Y, Tao Y, Zhang M, Wu D, Guan S, Han W, Ying W. Distinct Patterns of the Autofluorescence of Body Surface: Potential Novel Diagnostic Biomarkers for Stable Coronary Artery Disease and Myocardial Infarction. bioRxiv 330985, 2018.

15. Li Y, Zhang M, Tao Y, Ying W. Lipopolysaccharide (LPS) induces increased epidermal green autofluorescence of mouse. bioRxiv 501189, 2018.

